# An Arabidopsis gene expression predictor enables inference of transcriptional regulators

**DOI:** 10.1101/2020.04.07.029413

**Authors:** Haiying Geng, Meng Wang, Jiazhen Gong, Yupu Xu, Shisong Ma

**Affiliations:** Hefei National Laboratory for Physical Sciences at the Microscale, School of Life Sciences, Division of Life Sciences and Medicine, University of Science and Technology of China, Innovative Academy of Seed Design, Chinese Academy of Sciences, Hefei, China; School of Data Science, University of Science and Technology of China, Hefei, China

**Author notes:** These authors contribute equally (Haiying Geng, Meng Wang, Jiazhen Gong). Correspondence: Shisong Ma.

## Abstract

Gene expression regulation by transcription factors (TF) has long been studied, but no model exists yet that can accurately predict transcriptome profiles based on TF activities. We have constructed a universal predictor for Arabidopsis to predict the expression of 28192 non-TF genes using 1678 TFs. Applied to bulk RNA-Seq samples from diverse tissues, the predictor produced accurate predicted transcriptomes correlating well with actual expression, with average correlation coefficient of 0.986. Having recapitulated the quantitative relationships between TFs and target genes, the predictor further enabled downstream inference of TF regulators for genes and pathways, i.e. those involved in suberin, flavonoid, glucosinolate metabolism, lateral root, xylem, secondary cell wall development, and endoplasmic reticulum stress response. Our predictor provides an innovative approach to study transcriptional regulation.

## INTRODUCTION

Transcriptome, a critical determinant of cellular functions and status, is largely modulated by gene transcription. Transcription in turn is mainly managed by a network of transcription factors (TF) that integrate environmental and developmental cues to fine tune target genes’ expression (*1, 2*). Gene regulatory networks (GRN) that connect TFs to their target genes are key to decipher cellular functions and regulatory mechanisms (*3*). GRNs have been constructed systematically by characterizing TF binding motifs, by profiling TF-DNA interaction, or by perturbing TFs expression (*4-7*). They were also inferred from transcriptome data using computational approaches like probabilistic graphical model, neural network, LASSO regression, least angle regression, and decision tree (*8-12*). Nevertheless, for both plants and animals, comprehensive and universal models that quantitatively predict gene expression levels based on TF activities have yet to be built. Here, we describe an Arabidopsis gene expression predictor for reconstructing highly accurate transcriptome profiles based on TFs expression. The predictor is tissue-independent and can be universally applied to samples derived from all tissues. It also enables downstream inference of TF regulators for genes and pathways, thus facilitating mechanistic investigation on transcriptional regulation.

## RESULTS AND DISCUSSION

### A universal gene expression predictor for Arabidopsis

We aimed to construct a predictor that predicts Arabidopsis gene expression values based on TFs’ expression levels and use it to infer TF regulators for genes and pathways. A linear predictor model:

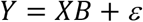

was built via ordinary least squares regression, where target genes’ expression matrix *Y* were estimated as linear combinations of TFs’ expression matrix *X*, with *B* and ε representing the coefficient matrix and random errors (Figure 1A). Large-scaled Arabidopsis RNA-Seq raw data were downloaded from NCBI’s Sequence Read Archive (SRA) database and processed via a uniform pipeline. After quality filtering, transcriptomes from 24545 RNA-Seq runs derived from diverse tissues (Figure 1B) were kept and merged into a large expression matrix for predictor model training. The predictor, after excluding low-expressed genes, shall take 1678 TF genes as inputs to predict the expression of 29182 non-TF genes.

**Figure 1.**
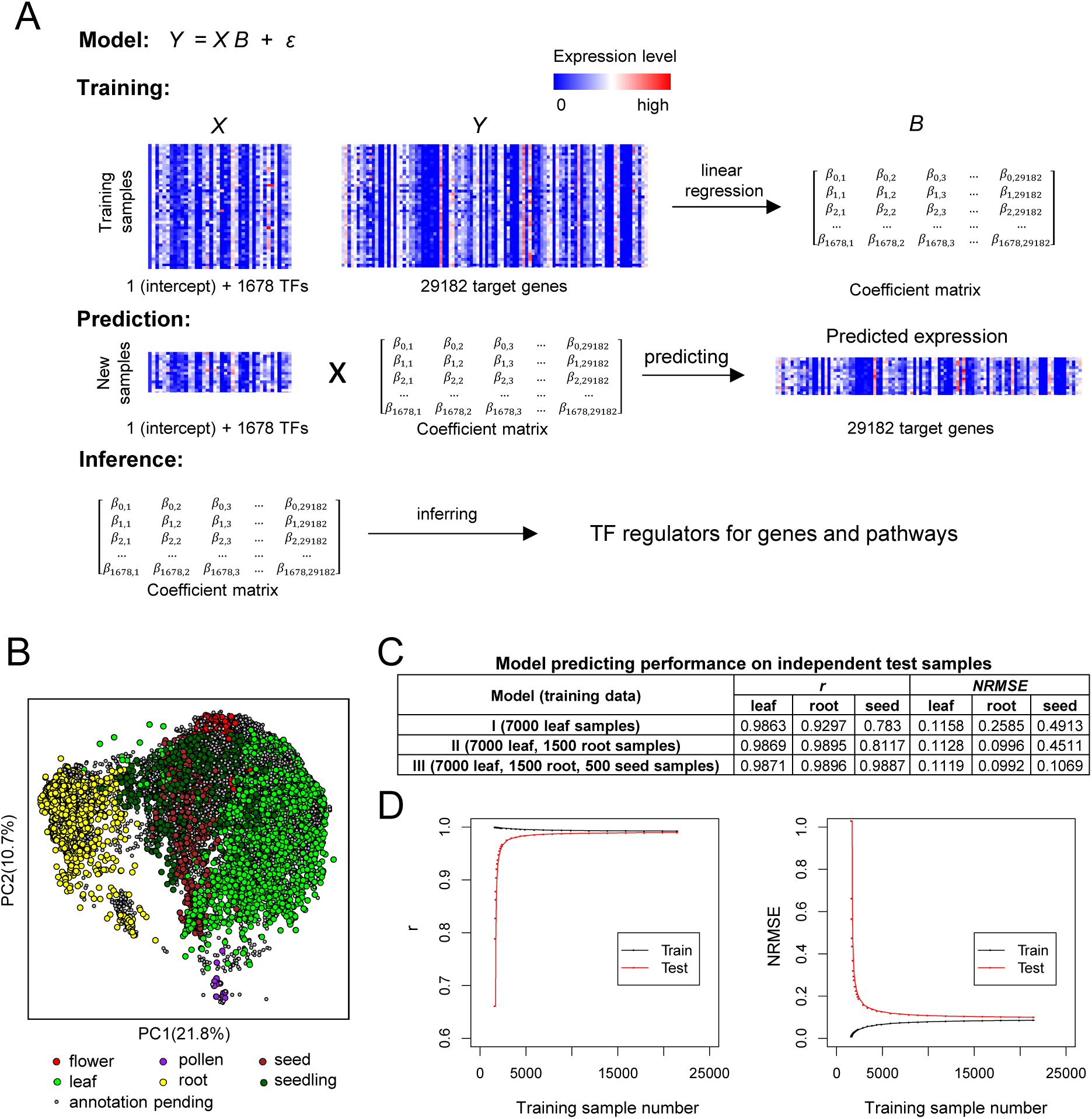
Building an Arabidopsis gene expression predictor. **(A)** The analysis workflow. **(B)** Principle component analysis of the 24545 RNA-Seq samples used for training the predictor model. PC, principle component. **(C)** Three predictor models trained with different combinations of leaf, root, and seed samples were tested for predicting performance on independent leaf (n=500), root (n=500), and seed (n=500) samples. Shown are the average Pearson’s correlation coefficients (*r*) and normalized root mean square errors (*NRMSE*) between the predicted and actual transcriptomes. **(D)** Predictor models trained with different number of training samples were tested for their performance. Shown are the average *r* (left) and *NRMSE* (right) between the predicted and actual transcriptomes for the training samples (black) and independent test samples (red, n=3000).

To obtain a universal predictor, the sample diversity and sample number of the training dataset are critical. A predictor model trained with leaf samples only performed poorly in predicting transcriptomes from roots and seeds, with low Pearson’s correlation coefficients (*r*) and high normalized root mean square errors (*NRMSE*) between the predicted and actual gene expression values (Figure 1C). Adding root and seed transcriptomes to the training dataset improved predicting quality for independent root and seed samples, while, interestingly, also improved prediction for independent leaf samples. Thus synergism was observed that combining transcriptomes from different tissues in the training dataset yielded better models than using single tissue alone. Since our model has a large number of input TFs, over-fitting might be a concern. A model trained with 1700 samples, despite possessing very high *r* (0.9999) for the training samples, had limited ability to predict independent test samples (with average test *r* 0.6612 and *NRSME* 1.0294), indicating high level of optimism or over-fitting within the model. As the number of training samples increased, the model’s ability to predict test samples improved, and the difference between the training and test *r* and *NRSME* decreased, suggesting the decline of over-fitting (Figure 1D). Indeed, a model trained with 20000 samples had average *r* of 0.9926 and 0.9896 for the training and test samples, respectively. The model explained 98.52% (*r*^*2*^) of the variances of gene expression values in the training samples, among which 0.6% might be due to over-fitting. Thus, increasing the number of training samples to 20000+ reduced over-fitting to a negligible level.

In light of the above results, all transcriptomes of the 24545 RNA-Seq runs were used to train the predictor model. To evaluate the predictor’s performance, RNAs were extracted from Arabidopsis root and shoot tissues to generate two RNA-Seq transcriptomes. Using the expression levels of 1678 TF genes, the predictor accurately predicted the expression of other 29182 non-TF genes for these two samples, with *r* > 0.994 and *NRSME* < 0.075 (Figure 2A). A leave-one-out cross-validation (LOOCV) strategy was also employed to assess the predictor’s ability to predict other novel independent samples. The 24545 RNA-Seq runs used for predictor training came from 1342 NCBI SRA studies. In each LOOCV run, a study was selected whose samples were held out, and the predictor was re-trained with all other RNA-Seq samples and then tested on the hold-out samples. This is similar to applying the predictor to novel independent samples. Figure 2B shows the LOOCV results for the study SRP075604 that had measured 138 transcriptomes across diverse Arabidopsis tissues (*13*). The predictor had highly accurate predictions for samples derived from all these tissues, with an average *r* of 0.9874, although the performance was slightly better for leaf and root samples than anther samples (Figure 2B). In total, LOOCV test was conducted for 546 studies that contain 10 or more RNA-Seq runs. The predictor had highly accurate predictions for samples from most of these studies, with average *r* of 0.9861±0.0177 and *NRSME* of 0.1066±0.0499 (Figure 2C). However, the predictor had moderate performance on single-cell RNA-Seq samples (data not shown). Single-cell RNA-Seq samples accounted for only a minor fraction of all Arabidopsis RNA-Seq datasets and they were excluded from our analysis. Thus, we have constructed a highly accurate and tissue-independent gene expression predictor that can be applied universally to most Arabidopsis bulk RNA-Seq samples.

**Figure 2.**
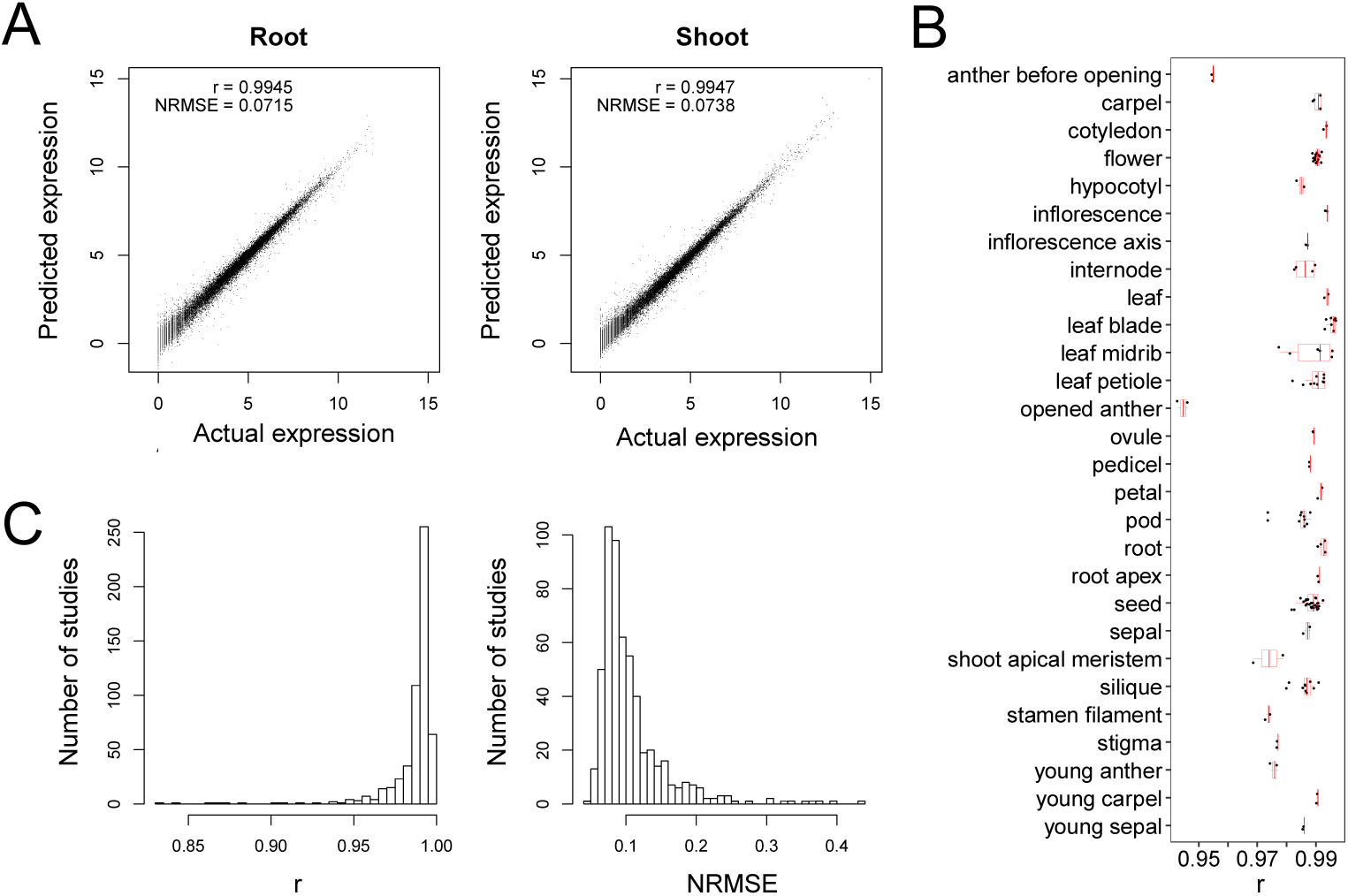
The predictor’s predicting performance on independent novel samples. **(A)** Two scatterplots showing the predicted and actual transcriptomes for two RNA-Seq samples derived from Arabidopsis root and shoot tissues. The two transcriptomes were generated in our lab to test the model’s performance. **(B)** A boxplot showing the leave-one-out cross-validation (LOOCV) test result for samples from the SRA study SRP075604 (*13*). **(C)** LOOVC test results for 546 SRA studies with 10+ samples. For each study, average LOOCV test performance statistics (*r* and *NRMSE*) were calculated for all its samples, which were then used to generate the histograms showing the performance distribution for all 546 studies.

### Arabidopsis gene co-expression modules

Besides the predictor model, gene co-expression modules were also identified using the same expression dataset. An Arabidopsis gene co-expression network based on the graphical Gaussian model (GGM) was constructed via a previously published procedure (*14*). The partial correlation coefficients (*pcor*) between all genes were calculated, and 312790 significant co-expressed gene pairs with *pcor* >= 0.045 were extracted to build a gene co-expression network (Figure S1 and Table S1). The network was then clustered into 1085 gene co-expression modules via the Markov Clustering (MCL) algorithm (Figure 3A and Table S2) (*15*). According to guilt-by-association, genes within the same module have similar expression patterns and might function in the same pathways. Indeed, 378 identified modules have enriched gene ontology (GO) terms (*P* <= 0.001) and function in pathways involved in, i.e., metabolism, development, and stress response (Table S3). For example, Module #84 was considered as a suberin biosynthesis module since it is enriched with 6 suberin biosynthetic genes (*P* = 3.82E-11), like *CYP86B1, ABCG20, ABCG6, GPAT5*, and *RWP1* (Figure 3B). Module #272 participates in glucosinolate biosynthesis (*P* = 9.04E-19) and contains key genes like *CYP79B3, SUR1*, and *UGT74B1*, while #210 is involved in flavonoid and anthocyanin biosynthesis (*P* = 1.02E-17) with genes like *TT3, TT18, At5MAT* (Figure 3C, D). Development related modules were identified as well, such as #105 for lateral root development (*P* = 1.86E-06), #65 for xylem development (*P* = 1.05E-14), and #11 for secondary cell wall biogenesis (*P* = 9.25E-43) in the late stage of xylem formation (Figure 3E-G). Also recovered were modules involved in stress response, such as #46 for endoplasmic reticulum (ER) stress response (*P* = 3.96E-50) (Figure 3H). The genes within these modules shared similar expression patterns and might be regulated by the same set of TFs. It is of great interest to identify such TFs as they could be the master transcriptional regulators for pathways that define a plant’s form and functions.

**Figure 3.**
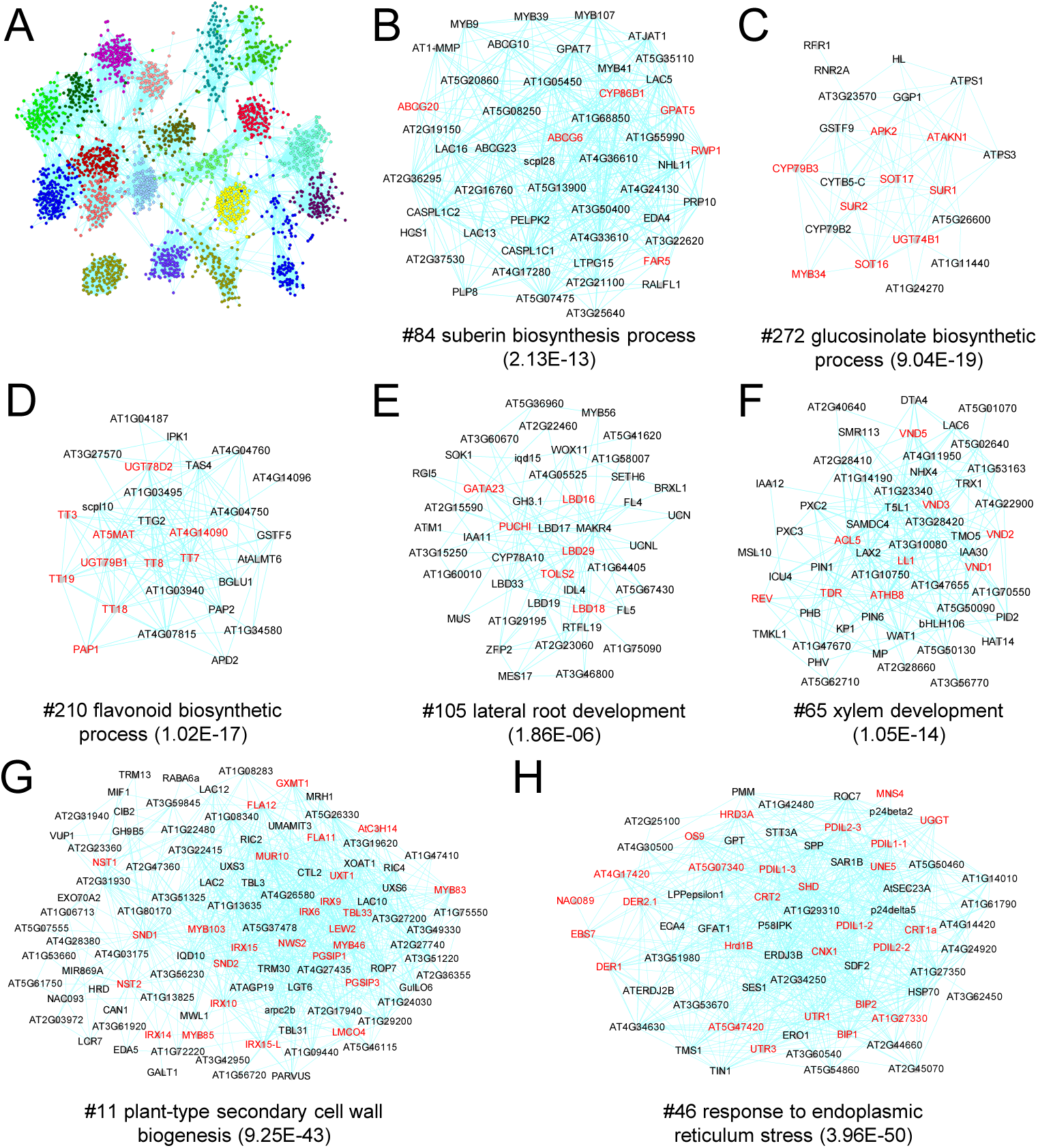
Gene co-expression modules identified via co-expression network analysis. **(A)** The 20 largest gene co-expression modules identified from the Arabidopsis GGM gene co-expression network. Dots represent genes, and co-expressed genes are linked by edges. Colors indicate module identities. Only the top 20 largest modules are shown due to space limit. **(B – H)** Subnetworks for selected gene co-expression modules that function in suberin (B), glucosinolate (C), flavonoid (D) biosynthesis, lateral root (E), xylem (F), secondary cell wall (G) development, and endoplasmic reticular stress response (H). The module id and represented enriched GO term (and its p-value) are shown for each module, with the genes possessing the represented GO term highlighted in red text.

### Transcriptional regulators revealed by the predictor

The predictor model and gene co-expression modules were then used to infer transcriptional regulators for genes and pathways. The predictor essentially recapitulated the quantitative relationships between TFs and target genes within its coefficient matrix *B*. Significance tests were conducted for individual regression coefficients β within the matrix, and 980736 significant interacting TF-target gene pairs corresponding to the coefficients with p-value <= 1E-09 (approximately equal to a Bonferroni-corrected p-value of 0.05) were extracted (Table S4). The TFs connected to a target gene by these significant TF-gene pairs were then considered as that gene’s predictor TFs. In average, each gene has 34 predictor TFs. These predictor TFs can be deemed as the gene’s potential transcriptional regulators. As an example, 37 TFs were identified as predictor TFs for *CYP86B1*, a gene encoding a fatty acid cytochrome P450 oxidase required for suberin biosynthesis (Figure 4A) (*16*). Its top predictor TFs include *MYB41, MYB107, MYB93, NAC058*, and *WRKY56*. Among them, *MYB41* and *MYB107* are transcriptional regulators of suberin biosynthesis, whose over-expression or mutation altered the expression of *CYP86B1*, while *MYB93*’s homologue from apple *MdMyb93* also modulates suberin deposition (*17-19*). Thus, among the 37 predictor TFs for *CYP86B1* include its known regulators and possibly other potential novel regulators as well.

**Figure 4.**
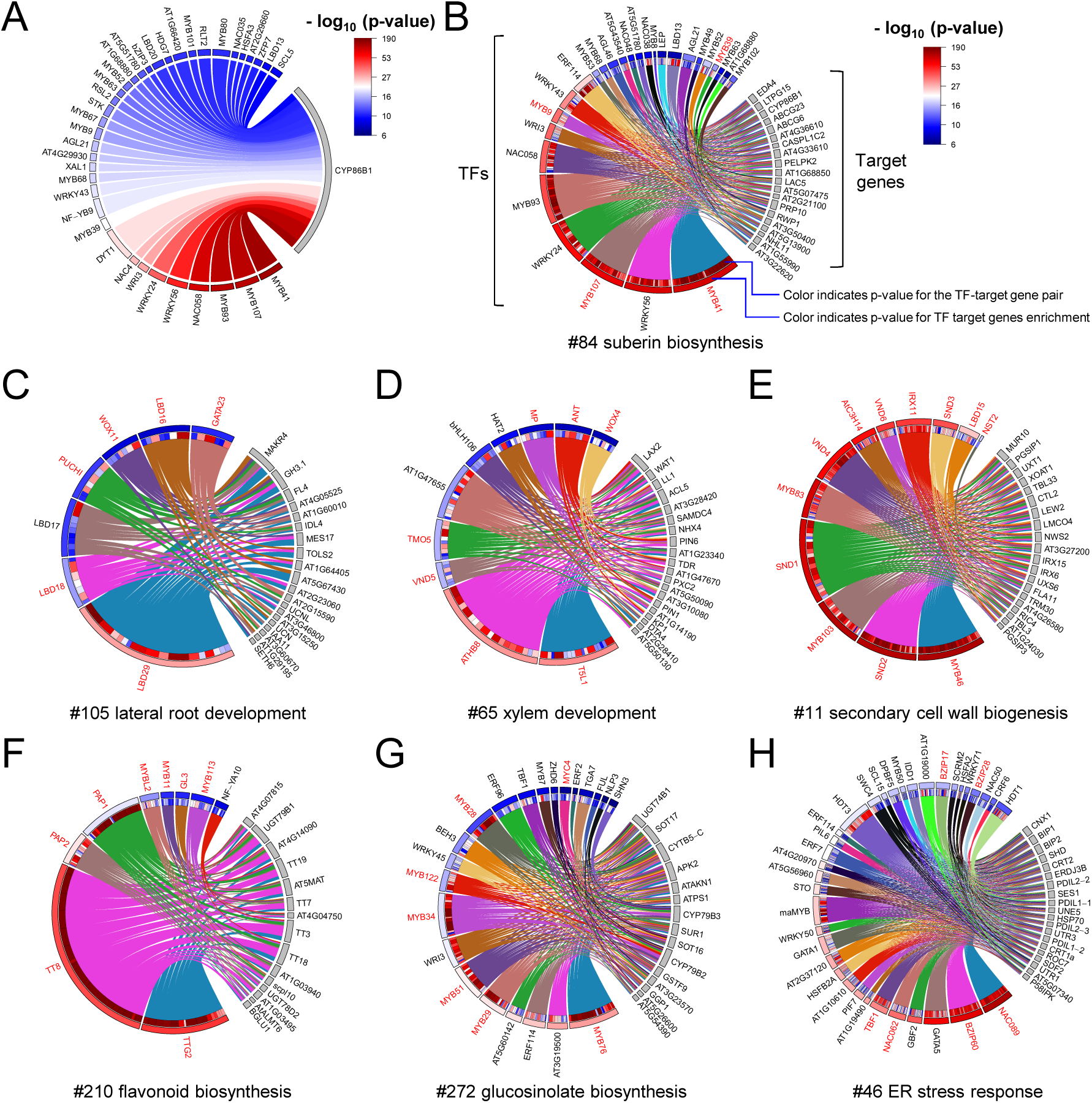
Predictor TFs identified for genes and pathways. **(A)** A Chord diagram linking the gene *CYP86B1* and its 37 predictor TFs. Colors indicate the p-value associated with the correspondent coefficients for the TF-target gene pairs extracted from the predictor, while the width of the links are proportional to the magnitude of the coefficient. **(B – H)** Chord diagrams linking the selected module’s predictor TFs to their target genes within the module. The links are color-coded by TF genes, their left ends’ colors indicate the p-values of the corresponding coefficients, while their widths are proportional to the coefficients’ magnitude. Also indicated are the p-values for the enrichment of the TF’s target genes within the module. Predictor TFs in red text are known regulators of the corresponding pathways. Due to space limit, only 15-20 target genes were shown.

Beyond individual genes, predictor TFs were also inferred at the pathways level for gene co-expression modules. As an example, Module #84 for suberin biosynthesis contains 45 non-TF genes, including *CYP86B1*. These genes’ common predictor TFs were then considered as the module’s predictor TFs. For instance, *MYB41* is shared by 42 non-TF genes within Module #84 as their predictor TF, while it has only 281 target genes among all non-TF genes. Thus *MYB41*’s target genes are enriched 97 folds within the module (*P* = 1.16E-82). A TF gene was considered as a predictor TF for a module if it was shared by at least 20% or 6, depending on which is larger, of the module’s non-TF genes as predictor TF and its target genes were also enriched within the module (*P* <= 1E-06). Under such criterion, 27 TFs, including *MYB41*, were identified as predictor TFs for Module #84, among which are known suberin regulators like *MYB41, MYB107, MYB9* (*17, 18*), and other MYB-, NAC-, and WRKY-type TFs as potential regulators (Figure 4B). Interestingly, among the potential regulators, *MYB39* was recently shown to be another regulator of suberization (*20*).

Predictor TFs were also identified for another 635 modules (Table S5). In average, 12 predictor TFs were identified for each module. Many known and potential novel TF regulators for diverse pathways were revealed. For example, predictor TFs were recovered for the development-related modules. Among them, Module #105, involved in lateral root development, had 7 predictor TFs, *LBD29, LBD18, LBD16, PUCHI, GATA23, LBD17*, and *WOX11*, all of which except *LBD17* have been shown to be regulators of lateral root development (Figure 4C) (*21, 22*). Module #65, for xylem development, had 10 predictor TFs, 7 of which are known modulators (*MP, WOX4, ATHB8, TMO5, T5L1, VND5, ANT*) of xylem or vascular development (*23, 24*) and the other 3 could be potential regulators (Figure 4D). On the other hand, all identified predictors for Module #11, functioning in secondary cell wall biogenesis, are known regulators of secondary cell wall biosynthesis, including *SND1, SND2, SND3, VND4, VND6, MYB46, MYB103, MYB83, IRX11, NST2, LBD15* and *AtC3H14* (Figure 4E) (*5, 25-28*).

Also revealed were predictor TFs for modules functioning in metabolism and stress responses. For example, Module #210, involved in flavonoid and anthocyanin biosynthesis, had 9 predictor TFs, including *TT8, TTG2, PAP1, PAP2, GL3, MYB11, MYB113, MYBL2* and *NF-YA10* (Figure 4F). Surprisingly, all of them except *NF-YA10* have been shown as regulators of flavonoid/anthocyanin biosynthesis (*29, 30*). In contrast, Module #272, participating in glucosinolate biosynthesis, had 22 predictor TFs, but only 7 of them (*MYB28, MYB29, MYB76, MYB34, MYB51, MYB122, MYC4*) have been shown to regulate glucosinolate biosynthesis (*31-33*), indicating there might be more potential regulators among the rest predictor TFs (Figure 4G). Similarly, the ER stress response module (#46) had 50 identified predictor TFs, among which 6 are known ER stress regulators (*bZIP60, bZIP17, bZIP28, NAC089, NAC062, TBF1*) and they cover most of the major TFs regulating ER stress that have been characterized so far (*34-36*), while the rest 44 could be potential modulators (Figure 4H).

Besides the examples mentioned above, modules and their predictor TFs were also identified for many other plant processes, such as the development of pollen tube (Module #1), anther (#78), Casparian strip (#138), root (#71), root hair (#5), flower (#132), shoot (#102), the metabolism of wax (#33), phenylpropanoid (#61), fatty acid (#47), nitrate (#15), phosphate (#38), sulfur (#91), iron (#58, 86, 222), and the response to heat (#125), cold (#170), water deprivation (#224), hypoxia (#35), and biotic stresses (#12) (Table S3 and S5). These examples demonstrated that our approach can effectively identify gene modules associated with a broad range of biological processes and their known and potential transcriptional regulator TFs.

Overall, we have constructed a highly accurate and tissue-independent gene expression predictor to predict Arabidopsis transcriptomes based on TFs expression. The key to obtain such a universal predictor was using a large number of transcriptomes derived from diverse tissues to train the predictor. Our results indicated the high-dimensioned gene expression profiles can be modeled as linear combinations of reduced-dimensioned TFs expression data, which is consistent with previous report that gene expression data have low dimensionality (*37*). There were also reports to use a small number of selected informative marker genes to impute gene expression profiles (*38, 39*). But these studies mainly focused on transcriptome prediction, without further mechanistic inference. In contrast, by using TFs as regressors to construct the model, our predictor not only achieved highly accurate predicted transcriptomes, but also recapitulated the quantitative relationships between TFs and target genes that seem to preserve across tissues, which further enabled downstream inference of TF regulators for genes and pathways. As demonstrated by the aforementioned examples, many of the identified predictor TFs for the modules are known regulators of the related pathways, while the rest could serve as novel potential regulators. The same approach, which we named as EXPLICIT (<u>Ex</u>pression <u>P</u>rediction via <u>Li</u>near <u>C</u>ombination of <u>T</u>ranscription Factors), also worked for other plant and animal species, such as rice, maize, mouse, and human (Wang et al. submitted, and unpublished data). Thus, for multiple eukaryotic species, there exist universal coefficient matrices that govern the quantitative relationships between TFs and target genes across tissues. These matrices provide the keys to understanding transcriptional regulation in biological systems.

## MATERIALS AND METHODS

### Gene expression data and gene expression predictor model

Raw data of publicly available Arabidopsis RNA-Seq runs were downloaded from NCBI SRA database. After removing single-cell or small RNAs sequencing runs and the runs with less than 1000000 reads, reads from the remaining runs were mapped against Arabidopsis genome (Araport11) using STAR 34 (version 020201) to produce read counts for each annotated genes (*40*). The cpm (counts per million) expression values were calculated for each gene, and the runs with unique mapping rates < 50% or total mapping rates < 70% or with < 5000 genes having cpm >= 1 were removed. Low-expressed genes that had cpm >= 1 in less than 100 samples were also filtered out. Finally, the expression values of 30860 genes in 24545 RNA-Seq runs from 1342 SRA studies were kept and merged into a large expression matrix after log transformation (log_2_(cpm+1)) for further analysis.

Using an Arabidopsis TF gene list obtained from PlantTFDB (*41*), two expression matrix for 1678 TF genes *X* and 29182 non-TF genes *Y* were extracted and used to train a gene expression predictor using a linear model *Y* = *X B* + *ε*, with *B* and *ε* representing coefficient matrix and random errors. The coefficient matrix was estimated in MATLAB (version R2019a) via least squares as *B = (X*^*′*^*X) \ (X*^′^*Y*). The significance of individual regression coefficients within the predictor model were also determined via t test, following hypothesis test method for multiple linear regression (*42*). The resulted p-values were used to extract significant interacting TF-target gene pairs from the predictor model. Predictor TFs for every non-TF gene were then identified as those TFs connected to the gene by these significant interacting TF-target gene pairs. A MATLAB script was developed in-house to conduct the analysis.

To test the predictor performance on transcriptome prediction, Arabidopsis RNA-Seq samples were processed via the same pipeline described above and two actual expression matrix for TFs *X*_*t*_ and non-TFs *Y*_*t*_ were extracted. A predicted expression matrix was estimated as *Y*_*p*_ *= X*_*t*_ *B. Y*_*p*_ and *Y*_*t*_ were then used to calculate the Pearson’s correlation coefficients *r* and normalized root mean squared errors (*NRMSE*) between the predicted and estimated transcriptomes. *NRMSE* for a given sample was calculated as:

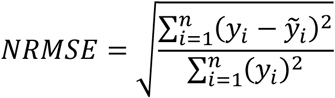

where *Y*_*i*_ and 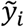 denote the actual and predicted expression values of gene i within the sample.

### Arabidopsis tissues and RNA-Seq sequencing

Surface-sterilized Arabidopsis seeds were stratified at 4°C in dark for 2 days and placed on vertical 1/2 MS Agar plates supplemented with 1% (w/v) sucrose and 0.8% (w/v) agar at 22 °C under 16h day/8h night cycle for growing. After 14 days, root and shoot tissues were harvested separately and total RNAs were extracted using TRI Reagent (Sigma-Aldrich, St. Louis, MO, USA). The quantity and integrity of the total RNAs were assessed on an Agilent 2100 Bioanalyzer (Agilent Technologies, Palo Alto, CA, USA), and only the samples with RIN values ≥ 8 were used for constructing mRNA sequencing libraries. mRNA-seq libraries were prepared for sequencing using NEBNext® Ultra™ RNA Library Prep Kit for Illumina® (New England BioLabs, Ipswich, MA, USA) according to manufacturer’s protocols. All libraries were sequenced using the Illumina NovaSeq platforms with paired-end 150 bp (PE 150) sequencing strategy. The library construction and sequencing were performed by the Novogene Corporation (Beijing, China).

### Gene module and predictor TF identification

Using the expression matrix composed of 30860 genes in 24545 RNA-Seq runs, an Arabidopsis gene co-expression network based on the graphical Gaussian model was constructed via a procedure published previously (*14*). Briefly, a random sampling strategy was used to calculate the *pcors* between all gene pairs. The procedure consisted of 20000 rounds. In each round, 2000 genes were randomly selected and *pcors* between them were calculated using the GeneNet package in R (*43*). After 20000 rounds, each gene pairs were sampled 83 times with 83 *pcors* calculated, and the *pcor* with lowest absolute value was then chosen as that gene pair’s final *pcor*. The gene pairs with *pcor* >= 0.045 were then chosen to construct a GGM gene co-expression network for Arabidopsis. The network was then clustered using the MCL algorithm (*15*), and 1085 gene modules that contained at least 6 genes were kept for further analysis. The network and its modules were visualized in Cytoscape (V3.4.0) (*44*). Arabidopsis GO terms annotations were obtained from TAIR (*45*), and GO enrichment analysis for the module were conducted using the hypergeometric distribution.

Predictor TFs were then identified for the modules. If a TF gene were shared by at least 6 as well as 20% of the non-TF genes within a module, and the TF’s target gene were also enriched within the module as determined via the hypergeometric distribution, the TF was considered as a target gene for the module. A perl script was developed in-house to identify predictor TFs for modules. The predictor TFs and target genes’ interactions within a module were visualized as Chord diagrams using the ‘circlize’ package in R (*46*).

## ACKNOWLEDGEMENT

This work was supported by grants from the National Natural Science Foundation of China (31770268), the Strategic Priority Research Program of the Chinese Academy of Sciences (XDA24010302), the Fundamental Research Funds for the Central Universities (WK2070000091), and University of Science and Technology of China (Start-up fund to S.M.). The numerical calculations in this manuscript were conducted on the supercomputing systems in USTC Supercomputing Center and USTC School of Life Sciences Bioinformatics Center.

## SUPPLEMENTAL FIGURES AND TABLES

**Figure S1**. Distribution of all partial correlation coefficients. A histogram showing the distribution of all *pcors* between 30860 genes. Only those gene pairs with *pcors* >= 0.045 were used for GGM gene co-expression network construction.

**Table S1**. Significant co-expression gene pairs used for GGM gene co-expression network construction.

**Table S2**. Module identities for the genes within the co-expression network.

**Table S3**. Gene Ontology enrichment analysis results for the modules.

**Table S4**. Significant interacting TF-target gene pairs identified form the predictor.

**Table S5**. Predictor TFs identified for the gene co-expression modules.

