## Supplementary figures and images for "An Arabidopsis gene expression predictor enables inference of transcriptional regulators"

### Figure S1

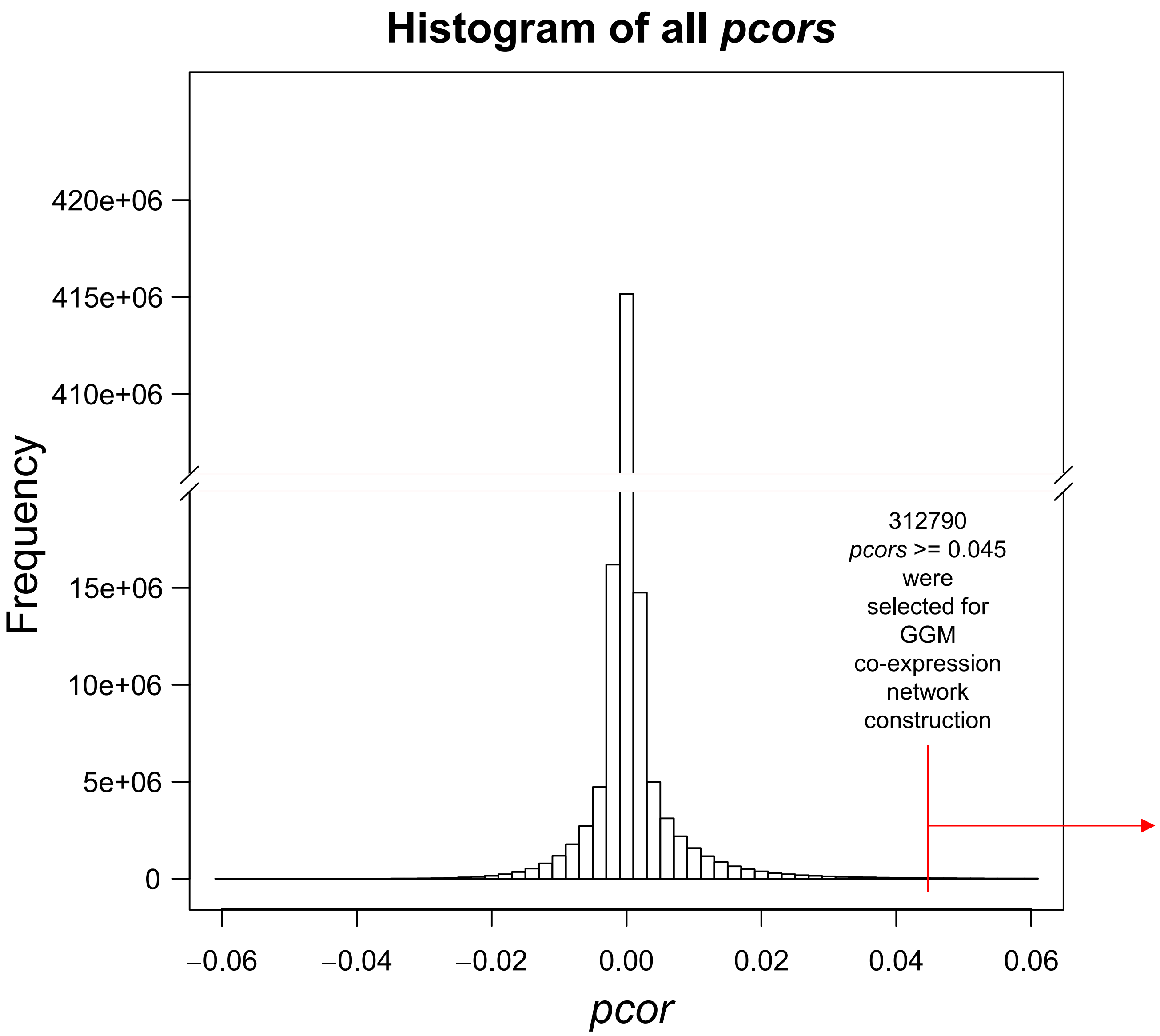
